# Oestradiol and Progesterone Variability in Adolescent Females: Associations with Brain Structure and Mental Health Problems

**DOI:** 10.64898/2025.12.28.696380

**Authors:** Muskan Khetan, Ye Ella Tian, Nandita Vijayakumar, Megan M. Herting, Michele O’ Connell, Marc L Seal, Sarah Whittle

## Abstract

**Objective:** Adolescent females are at increased risk for adverse mental health problems, and puberty is thought to be a critical factor. Pubertal onset involves substantial variability in oestradiol (E2) and progesterone (P4), yet the impact of these hormone dynamics remains unclear. We investigated whether within-individual E2 and P4 variability across one month relates to brain structure and mental health problems in adolescent females.

**Methods:** Participants were 147 females (11–17 years) from the community-based, cross-sectional Puberty and NeuroDevelopment in Adolescents (PANDA) study in Australia. Salivary E2 and P4 were collected weekly for one month, with variability indexed as within-subject standard deviation. Structural MRI was acquired on the final sampling day. Mental health problems were assessed via self- and parent-report. Linear regression and mediation analyses tested associations between hormone variability, brain structure, and mental health, adjusting for demographics.

**Results:** Greater P4 variability was associated with smaller left thalamus volume (Cohen’s d = −0.26, pFDR = 0.031). Higher E2 variability was associated with fewer anxiety (d = −0.170, pFDR = 0.043) and depressive symptoms (d = −0.179, pFDR = 0.036), lower emotion dysregulation (d = −0.179, pFDR = 0.036), and smaller right hippocampus volume in pre-menarchal adolescents (d = −0.192, p = 0.02). Weekly reductions in E2 predicted lower positive affect (d = 0.084, p = 0.048).

**Conclusion:** These findings indicate that hormone variability may meaningfully shape brain and behavioural development in adolescent girls differently from hormone-related transitions in adulthood. Larger, replicable, and longer-term studies are needed to confirm these patterns.

Oestradiol and Progesterone Variability in Adolescent Females: Associations with Brain Structure and Mental Health Problems. Muskan Khetan 1, Ye Ella Tian 2, Nandita Vijayakumar 3,4, Sarah Whittle 1,4

1. Centre for Youth Mental Health, University of Melbourne, Australia
2. Department of Psychiatry, University of Melbourne, Australia
3. Murdoch Children’s Research Institute, Australia
4. School of Psychology, Deakin University, Australia

## 1. Introduction

Adolescent females have a higher vulnerability to experiencing emotional dysregulation and a range of mental health problems than adolescent males^1,2^. Pubertal factors may contribute to this risk by influencing brain maturation^3^. During female puberty, oestradiol (E2) and progesterone (P4) begin to cycle across the month, but this hormonal cycling is marked by high variability, i.e., variability in the hormonal cyclical patterns across the month, comparable to other transitional stages such as peri-menopause^4^. While research has examined how hormone variability during postpartum and perimenopause affects brain structure^5^, far less is known about the adolescent period. Investigating how hormone variability influences adolescent brain structure could offer crucial insights into why females are at increased risk for many mental health problems during this stage of development.

During adolescence, female ovarian E2 and P4 rise and begin cycling during puberty, marking the onset of the reproductive stage^6^. Both E2 and P4 follow a distinct cyclical (or pulsatile) pattern throughout the menstrual cycle. E2 peaks twice, first in the late follicular phase (just before ovulation) and again alongside P4 during the mid-luteal phase (around seven days before menstruation). Both hormones decline during the premenstrual phase (or luteal phase)^7^. Notably, adolescent menstrual cycles are often irregular in this pattern, resulting in greater individual variability in hormonal patterns, i.e., a greater rise and fall of these hormones in a non-cyclical (non-pulsatile) manner, compared to adults^11^.

Although limited, existing evidence suggests that adolescents are sensitive to hormone variability, and this may begin even before menarche. For instance, greater E2 variability is associated with increased moodiness within a day in premenarchal girls as young as 9–10 years old^12^. Similarly, weekly changes in estrone (a urinary metabolite of E2) have been associated with greater mood sensitivity in girls aged 11–14, with stronger effects observed in premenarchal girls^4^. Menstrual cycle-related mood disorders such as premenstrual syndrome (PMS) and premenstrual dysphoric disorder (PMDD) are also common in adolescent females^13^, and are characterised by mood disturbances, anger/irritability, reduced interest in usual activities, and emotion dysregulation during the luteal phase of the menstrual cycle^14^. Similar behavioural patterns are observed during other hormonally dynamic life stages, such as post-pregnancy and perimenopause, where greater variability in E2 and P4 has been linked to higher levels of depressive symptoms^15,16^. Together, these findings suggest that hormonal variability may contribute to individual differences in emotion dysregulation and vulnerability to mental health difficulties across key transitional periods in the female lifespan. However, evidence during adolescence remains limited, and the underlying neural mechanisms are not yet well understood.

One possible mechanism by which hormonal variability is associated with vulnerability to mental illness symptoms and emotion dysregulation could involve the effects of hormones on brain structure during this critical period of neurodevelopment^17^. During puberty, E2 and P4 have key organisational effects on the brain via membrane-associated and intracellular receptors^18,19^. These hormone receptors are concentrated in regions that are involved in emotion regulation and are implicated in anxiety, depression and emotion dysregulation, such as the prefrontal cortex (PFC), amygdala, and hippocampus^20–22^. Prior adolescent studies have reported positive and negative associations between E2 levels and gray matter (GM) structure in PFC, parietal, middle temporal region and amygdala, and between P4 levels and GM structure within the default mode network that includes medial PFC, posterior cingulate cortex, medial and lateral temporal regions^23,24^. However, these studies have typically relied on average hormone levels, which may mask the impact of intra-individual variability. Fluctuations in hormone levels (or non-cyclical changes in hormones) within an individual during this sensitive developmental window may uniquely exert some influence on brain structure.

To date, only one study, to our knowledge, has investigated links between E2 variability and brain structure and mental illness symptoms in adolescent females^25^. This study found a negative association between individual E2 variability and medial orbitofrontal cortical thickness; however, there was no association with internalising symptoms in this relatively small sample (n = 44). Notably, no adolescent study, to our knowledge, has explored the effects of P4 variability on brain structure or mental illness symptoms. In adult studies, women with menstrual cycle related mood disorders show structural alterations in brain regions rich in E2 and P4 receptors—such as the hippocampus, parahippocampus, and amygdala^26,27^. Further, alterations in amygdala structure were also negatively associated with depressive symptoms in this population^26^. These findings underscore the importance of understanding how individual differences in hormone variability may shape brain structure (particularly the regions involved in emotion regulation) and mental illness symptoms in a larger adolescent sample.

Thus, the current study aimed to investigate how individual variability in E2 and P4 levels across a month relates to GM structure, focusing on brain regions involved in emotion regulation—the PFC, hippocampus, and amygdala^28^. Given the widespread effects of average E2 levels across the brain^29^, we also employed an exploratory whole-brain approach. We further aimed to investigate associations between hormone variability and internalising and externalising symptoms and emotion regulation difficulties. Links with internalising and externalising symptoms were investigated, given higher rates of these symptoms in females with menstrual cycle related mood disorders^30^. Finally, we aimed to test whether individual differences in brain structure mediate links between E2/P4 variability and mental health problems (defined as mental illness symptoms and emotion dysregulation in this study).

We hypothesised that variability in E2 and P4 would be associated with the structure of brain regions involved in emotion regulation, given that these regions are dense in E2 and P4 receptors^31^. Further, we hypothesised that there would be a significant association between E2/P4 variability and mental health problems via these brain structures. Due to the lack of adolescent research, we do not specify directional hypotheses.

## 2. Materials and Methods

Study hypotheses and analyses were preregistered (**10.17605/OSF.IO/RB84H**). Non-preregistered analyses are noted as exploratory.

### 2.1 Participants

Female participants aged 11 to 17 years were drawn from the Puberty and NeuroDevelopment in Adolescents (PANDA) Study. The PANDA Study is a cross-sectional study that recruited participants from Melbourne, Australia. Ethics approval was granted through the Royal Children’s Hospital Human Ethics Research Office (#1955580). Participants diagnosed with any developmental or endocrine disorders were excluded from this study. Additionally, those taking hormonal contraceptives were also excluded (for detailed inclusion and exclusion criteria, refer to the study protocol)^32^. A subset of individuals with clinical and MRI assessments, and weekly saliva samples over one month (i.e., minimum four saliva samples, more details in SM), were used in the current study (n=145, Figure 1). Data quality control and outlier detection are provided in the Supplementary Material (SM).

**Figure 1:**
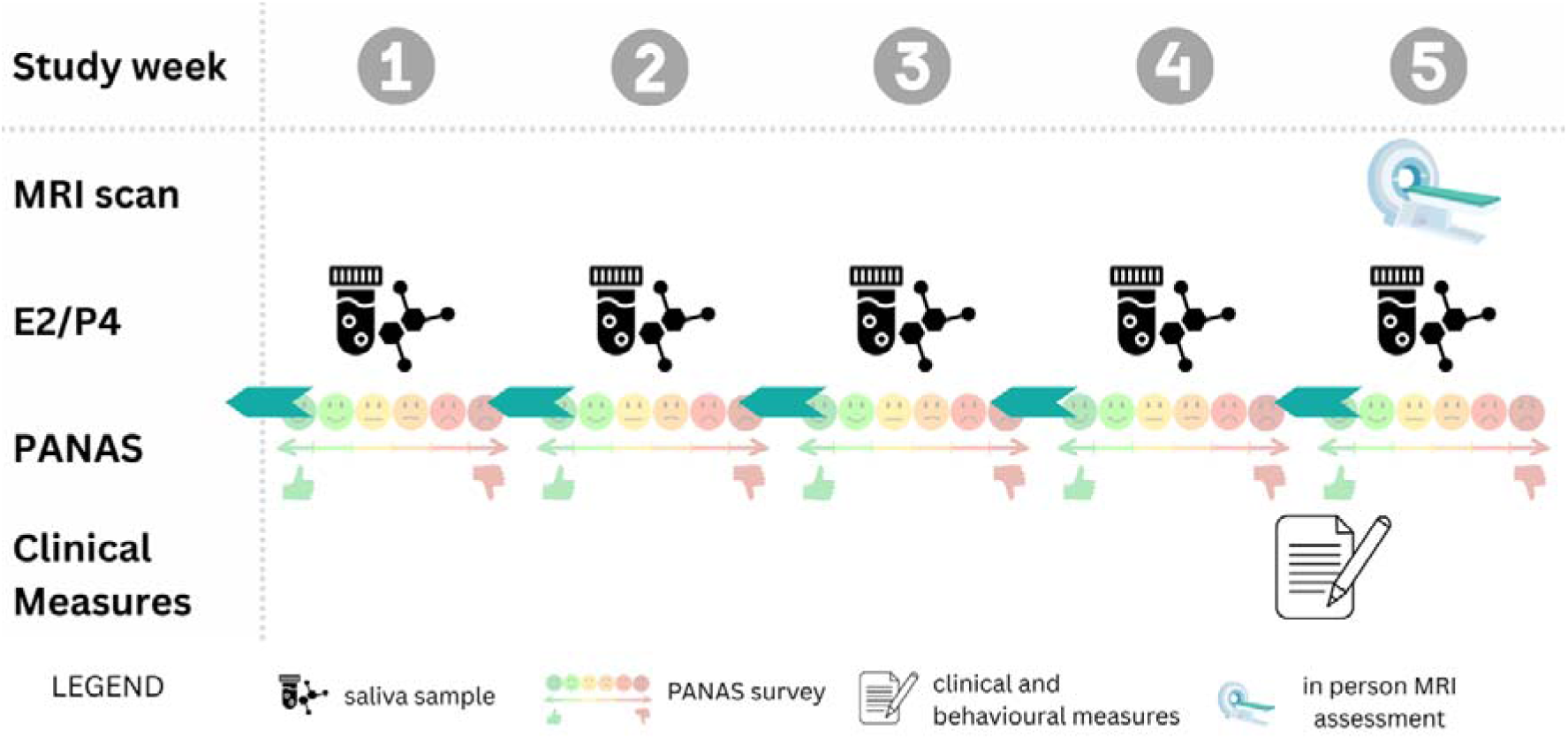
Study overview. Each week, participants collected saliva samples every morning to assay oestradiol (E2) and progesterone (P4). Along with saliva samples, each participant recorded their mood in the past week using the Positive and Negative Affect Schedule (PANAS). Clinical measures, including mental health symptoms and emotion regulation, were collected during the week before the in-person MRI visit.

### 2.2 Hormone (E2 and P4) acquisition and processing

Participants collected saliva samples immediately after waking (before eating, drinking, or brushing their teeth) once per week for four weeks before their MRI scan, plus an additional sample on the day of the scan (five samples total). Each sample (∼2.5 mL) was collected via passive drool into 10 mL Techno Plas centrifuge tubes using a straw. After collection, samples were frozen in participants’ homes within a sealed Bio-Bottle® container until the day of their MRI assessment and transported on ice to the assessment site. Upon arrival, samples were stored at −30°C until processing. Saliva was then thawed, centrifuged, and the supernatant was assayed for estradiol (E2) and progesterone (P4) by the Vaccine Immunology Group at MCRI (Murdoch Children’s Research Institute) using Salimetrics ELISA kits, with an E2 assay range of 1 – 32 pg/ml and a P4 assay range of 10 – 2430 pg/ml.

#### E2 & P4 variability measures

We indexed intra-individual E2 and P4 variability using the standard deviation (SD) of each participant’s hormone measurements, consistent with prior adolescent work^25^. Although alternative metrics such as the mean absolute successive difference (MASD) and mean squared successive difference (MSSD) have been proposed^4^, these were highly correlated with SD in our sample (see SM). We therefore used SD as the primary variability measure.

### 2.3 Magnetic resonance imaging (MRI)

#### Data acquisition

Neuroimaging data were acquired on the 3T Siemens MAGNETOM Prisma fit scanner – 32 channel head coils (Siemens, Erlangen, Germany) at the Royal Children’s Hospital, Melbourne, Australia. Due to software upgrades, some sequence parameters were different in subsets of participants. However, the majority of the T1-weighted scans acquired (∼80%) had the following parameters: repetition time = 2500ms; echo time1 = 1.72ms; echo time2 = 3.45ms; echo time3 = 5.18ms; echo time 4 = 6.91ms), flip angle = 8°; field of view = 256 mm; 208 contiguous 0.9 mm thick slices (voxel dimensions = 0.9mm^3^). Details on the imaging parameters for other participants are in SM.

#### Data preprocessing

T1-weighted images were processed using FreeSurfer v7.0.0 (http://surfer.nmr.mgh.harvard.edu/). Cortical reconstruction and segmentation were done with the recon-all function from FreeSurfer with 10 mm FWHM smoothing. This pipeline includes intensity normalisation, skull stripping, surface reconstruction, and segmentation, yielding vertex-wise cortical thickness and surface area measures^33,34^. Details on T1-weighted quality control procedures are in SM Subcortical segmentation was based on the *aseg* atlas^35^, providing volumes of 40 regions (20 for each hemisphere), of which we included 14: thalamus, caudate, putamen, pallidum, hippocampus, amygdala, and nucleus accumbens for each hemisphere.

### 2.4 Mental illness symptoms and Emotion regulation

Internalising symptoms were measured using the self-report Spence Children’s Anxiety Scale-Short (SCAS-S)^36^ and Children’s Depression Inventory 2 (CDI-2)^37^. Externalising symptoms were measured using the parent-report Strengths and Difficulties Questionnaire (SDQ)^38^. Emotion regulation was measured using the self-report Difficulties in Emotion Regulation Scale – Short Form (DERS-SF)^39^. All scales demonstrated good internal consistency in the current sample (Cronbach’s α = 0.72–0.90; see SM for details).

### 2.5 PANAS

Positive and negative affect over the past week was measured using the Positive and Negative Affect Schedule (PANAS)^40^. The PANAS has demonstrated good validity and internal consistency, with Cronbach’s α = 0.87 for positive affect and Cronbach’s α = 0.86 for negative affect in our sample.

### 2.6 Covariates

The current study included both pre- and post-menarchal adolescents. Menarche status was assessed using the self-report Pubertal Development Questionnaire (PDQ)^41^, which asked participants whether they had begun menstruating and their age at first period. Given that we were interested in the effects of E2 and P4 variability over and above pubertal development, menarche status was included as a categorical covariate (yes/no) in all analyses.

Sociodemographic covariates included race (caregiver-reported), caregiver’s education, and income-to-needs ratio (derived from after-tax household income and number of children). Given that the participants were scanned using different MRI sequences, we controlled for this as a dummy variable.

### 2.7 Statistical analyses

There were some missing values for CDI (N = 4), DERS-SF (N = 10), SCAS (N = 1) and income-to-needs ratio data (N = 6). All missing values for each variable were handled using multiple imputation (“mice” package in R), with 50 iterations. We then used the “merge_imputations” function^42^ in R to impute the missing values with the mean imputed value across all the iterations.

Mental illness symptoms and emotion regulation scores were transformed using the “transformTukey” function (R package *rcompanion*) to address skewness in the raw data. Additionally, normality of residuals was checked for all models, and if there were any deviations from normality, robust linear regression (“lmrob”) was used. Finally, influential outliers were checked using the Cook’s distance.

#### Association between hormone variability and brain structure

For the hypothesis-driven analysis, we used a PFC mask comprising cortical labels defined by the Desikan-Killiani atlas^34,35^, which included merging the medial orbitofrontal cortex, lateral orbitofrontal cortex, superior frontal, rostral, and caudal middle frontal cortex, pars orbitalis, pars triangularis, pars opercularis, rostral and caudal anterior cingulate labels for the left and right hemispheres separately.

We performed both whole-brain and masked vertex-wise analyses using Freesurfer’s general linear model analysis (“mri_glmfit”). In each model, E2 or P4 variability was entered as the predictor, with structural brain measures—cortical thickness and surface area at each vertex across the whole brain, and within the PFC mask—as the outcomes. To control for individual differences in brain size, surface area was scaled using estimated total intracranial volume (eTIV) during preprocessing. Vertex-wise results were corrected for multiple comparisons using Monte Carlo cluster-wise correction (cluster-forming threshold p < 0.001; FWE-corrected threshold p < 0.05)

For the subcortical volume analysis, the same linear regression model was fitted in R (using “glm” package), with volume at each subcortical region as an outcome and eTIV as an additional covariate in the models. To control for Type I error, false discovery rate (FDR) correction was applied across the 14 subcortical regions, with corrected p-values < 0.05 considered statistically significant.

#### Association between hormone variability and mental health/emotion regulation

Linear regression models were run in R (using the “glm” function) to examine the relationship between hormone variability and the outcomes. Mental illness symptoms (SCAS, CDI, SDQ) and emotion dysregulation (DERS) were entered as separate outcome variables. For any significant associations, follow-up analyses were conducted on relevant subscales. To adjust for multiple comparisons, FDR correction (p < 0.05) was applied.

#### Mediation

For any significant cortical clusters or subcortical volumes detected in the aforementioned analyses, mediation models were tested with mental health problem measures as outcomes, E2 or P4 variability as the predictor, and brain measures as the mediator. The mediation model was fitted using the R package “mediation” with a bootstrapping method (boot = 5000).

### 2.8 Exploratory analyses

To test whether any significant findings were specific to hormone variability, in exploratory analyses, we tested mean hormone values as predictors. We also tested both menarchal status and cycle irregularity (pre-menarche, regular, and irregular cycling) as moderators. We additionally examined whether week-to-week changes in hormone levels were associated with weekly mood (positive and negative affect) changes. Refer to SM for further details on the classification of cycle irregularity.

## 3. Results

Demographic, pubertal and clinical characteristics (DERS-SF, CDI, SCAS, and SDQ) of the final sample are shown in Table 1.

**Table 1:**
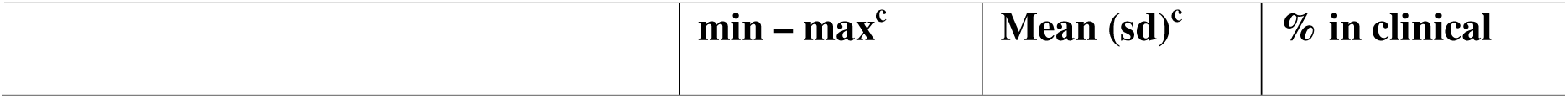
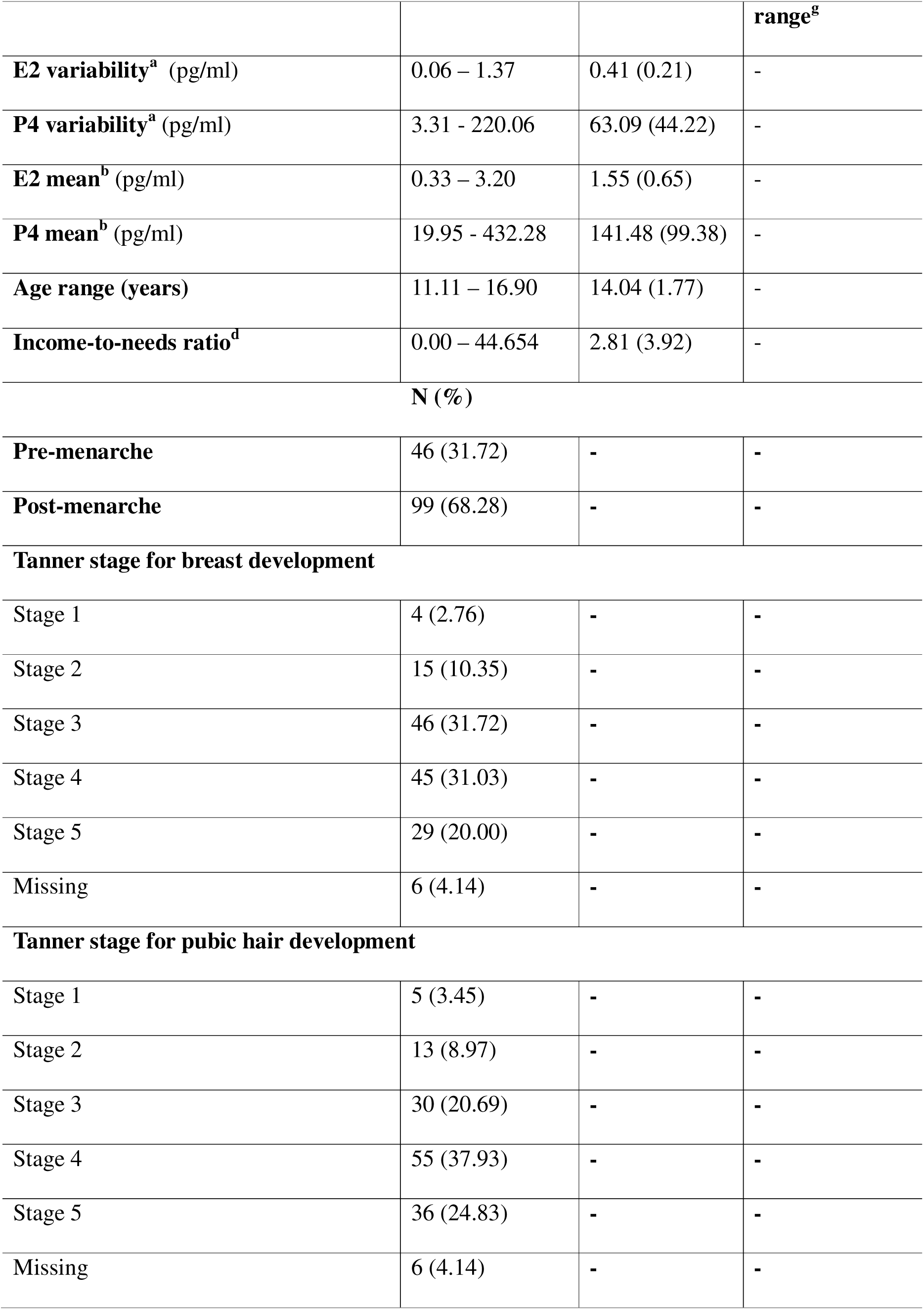
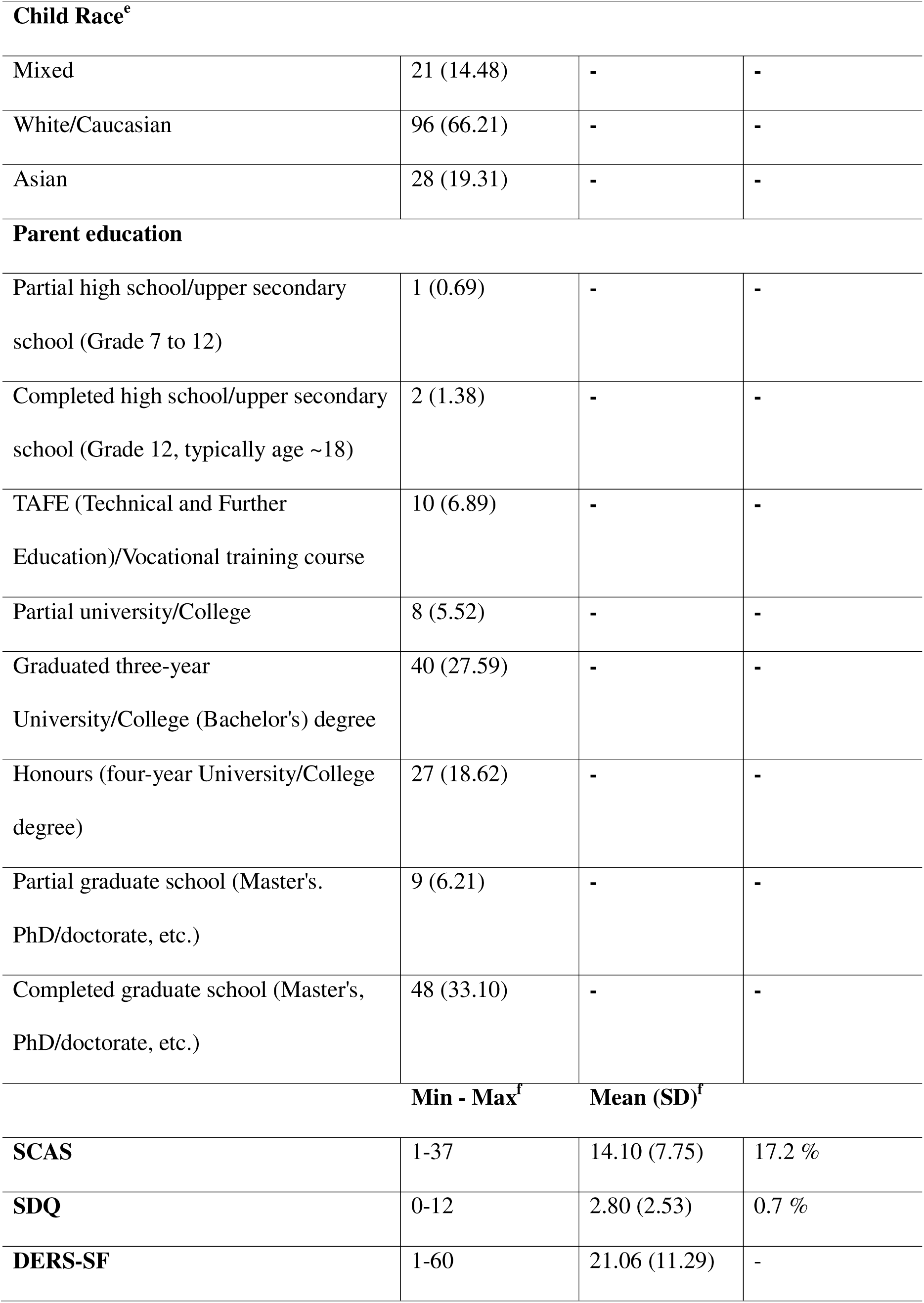
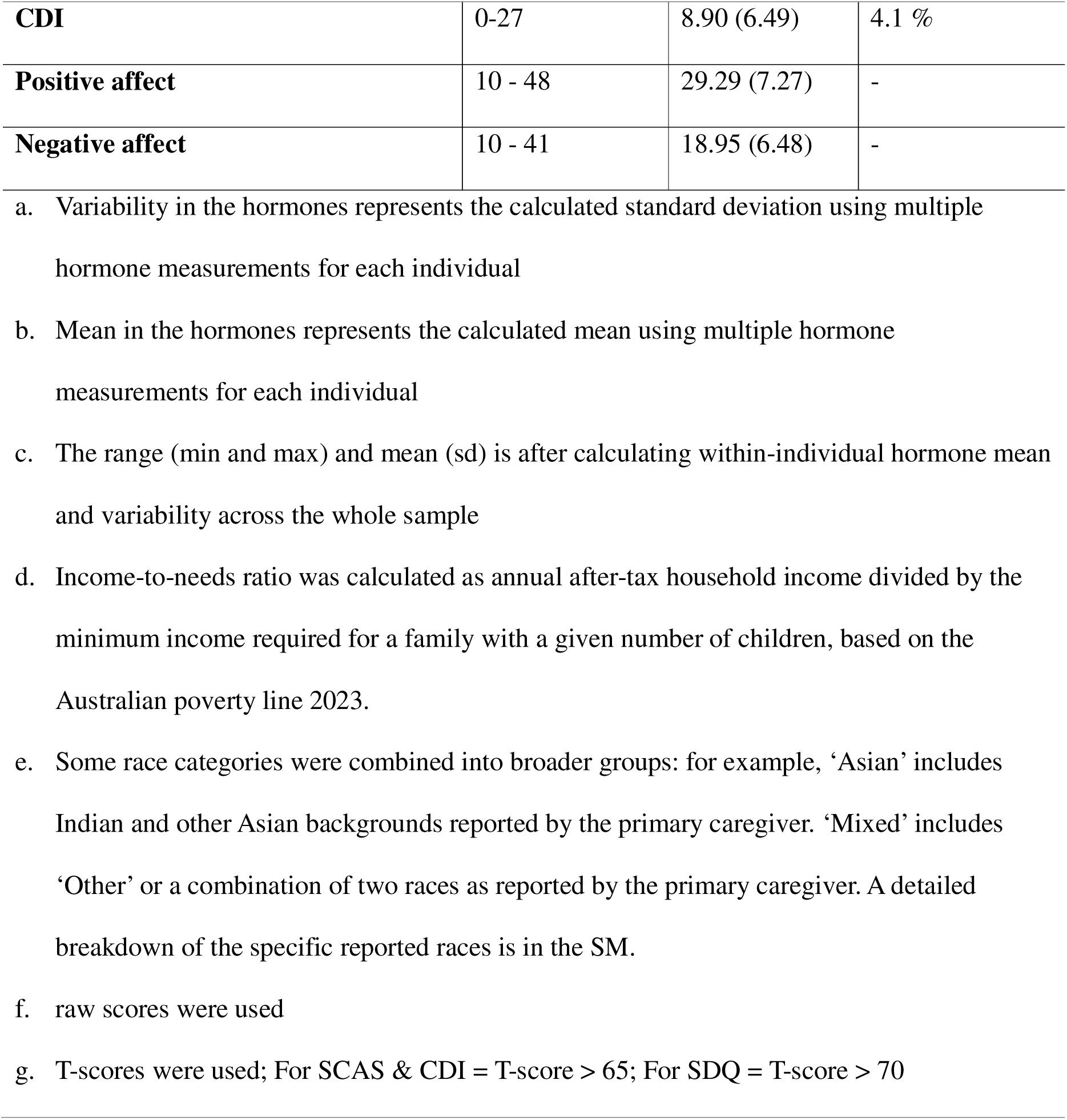
Descriptives of the final sample.

### 3.1 Association between hormone variability and brain structure

There were no significant association between hormone variability and cortical structure using vertex-wise analysis (refer to uncorrected cortical images in the repository https://neurovault.org/collections/IVDQQFDM/) Linear regression models showed a significant negative association between P4 variability and the volume of the left thalamus (Cohen’s d = −0.26, pFDR = 0.031; refer to Figure 2). There were no other significant associations between hormone variability and subcortical structure (see Table 2 for subcortical results).

**Figure 2:**
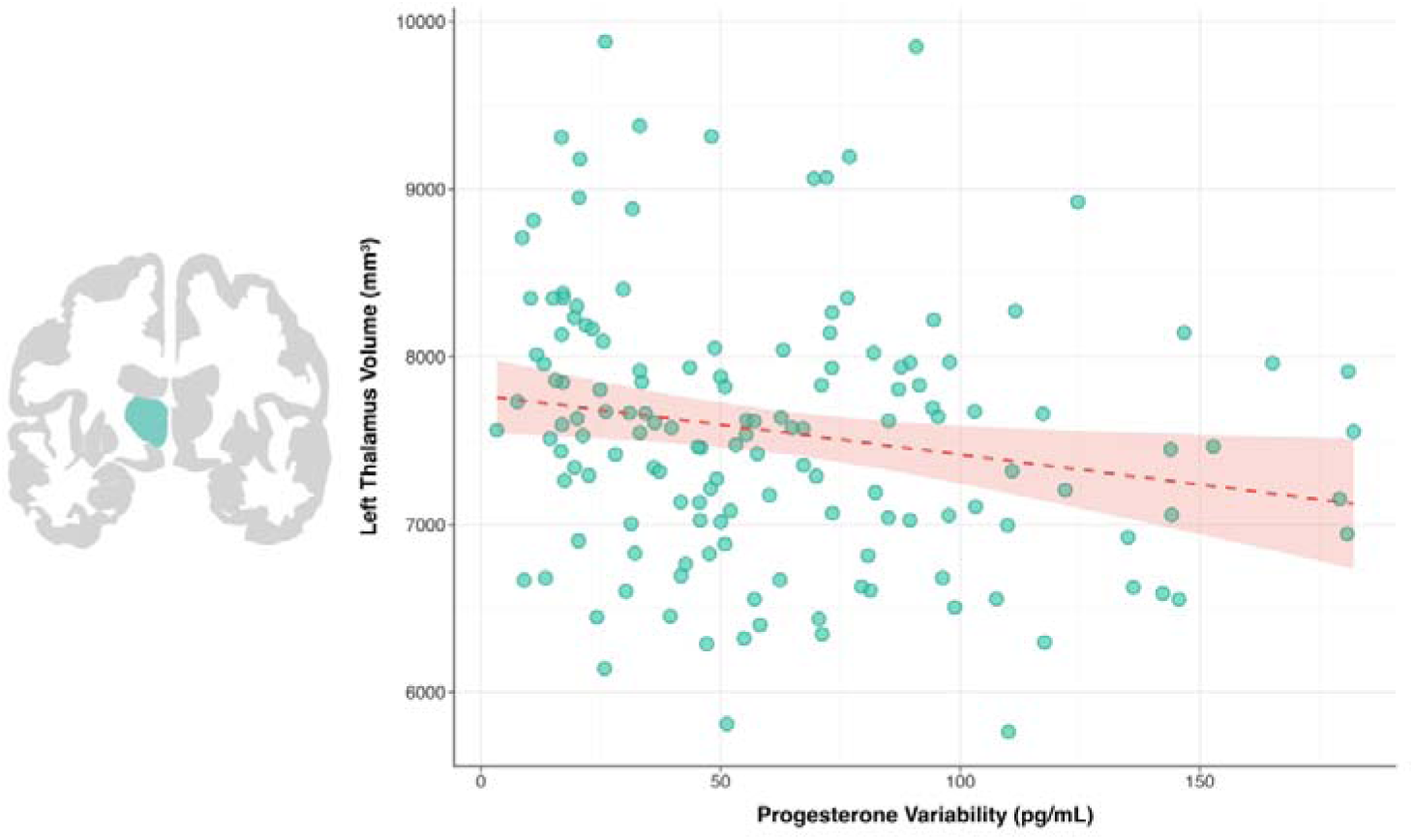
The fitted dashed line with 95% CI (shaded area) shows a negative association between progesterone variability (pg/ml) and volume in the left thalamus (mm^3^). Linear models were adjusted for menarche status, age at the time of MRI, adolescent’s race, primary caregiver’s education, income-to-needs ratio, a dummy variable for different T1-w sequences, and ICV

**Table 2:** Linear model between subcortical regions and hormone variability^a^.

| Subcortical regions (mm<sup>3</sup>) | P4 variability (pg/ml) |  | E2 variability (pg/ml) |  |
| --- | --- | --- | --- | --- |
|  | Cohen's D | pFDR | Cohen's D | pFDR |
| Left Thalamus | -0.259 | 0.031<sup>b</sup> | -0.168 | 0.435 |
| Right Thalamus | -0.133 | 0.775 | -0.103 | 0.720 |
| Left Caudate | 0.022 | 0.856 | 0.079 | 0.720 |
| Right Caudate | -0.032 | 0.856 | 0.060 | 0.720 |
| Left Putamen | -0.024 | 0.856 | -0.054 | 0.720 |
| Right Putamen | 0.010 | 0.903 | -0.060 | 0.720 |
| Left Pallidum | -0.046 | 0.814 | -0.025 | 0.770 |
| Right Pallidum | -0.077 | 0.814 | -0.042 | 0.770 |
| Left Hippocampus | -0.077 | 0.814 | 0.024 | 0.770 |
| Right Hippocampus | -0.075 | 0.814 | -0.065 | 0.720 |
| Left Amygdala | -0.057 | 0.814 | -0.103 | 0.720 |
| Right Amygdala | -0.050 | 0.814 | -0.157 | 0.435 |
| Left Accumbens | -0.064 | 0.814 | 0.031 | 0.770 |
| Right Accumbens | 0.079 | 0.814 | 0.062 | 0.720 |

- a. Linear models were adjusted for menarche status, age at the time of MRI, adolescent's race, primary caregiver's education, income-to-needs ratio, a dummy variable for different T1-w sequences, and ICV - b. Significant after FDR correction

### 3.2 Association between hormone variability and mental illness symptoms/emotion regulation

Hormone variability was not significantly associated with mental illness symptom/emotion regulation (p > 0.05), either before or after FDR correction. However, after checking for influential outliers using the Cook’s distance method, two influential outliers were found for models testing associations between E2 variability and symptoms (see SM, Figure 1 for Cook’s distance statistics and model output). After removing these outliers, there was a significant negative association between E2 variability and DERS-SF (d = −0.179; pFDR = 0.036), CDI (d = −0.179; pFDR = 0.036) and SCAS total (d = −0.170; pFDR = 0.043) scores (as shown in Figure 3 and Table 3). Follow-up models tested associations between E2 variability and DERS-SF, CDI and SCAS sub-scales. The models showed significant negative associations between E2 variability and DERS ‘strategies’ (d = −0.243; pFDR = 0.010), CDI ‘emotional problems’ (d = −0.208; pFDR = 0.028), and SCAS ‘separation’ (d = −0.302; pFDR = 0.001) and ‘social anxiety’ (d = 0.226; pFDR = 0.007).

**Figure 3:**
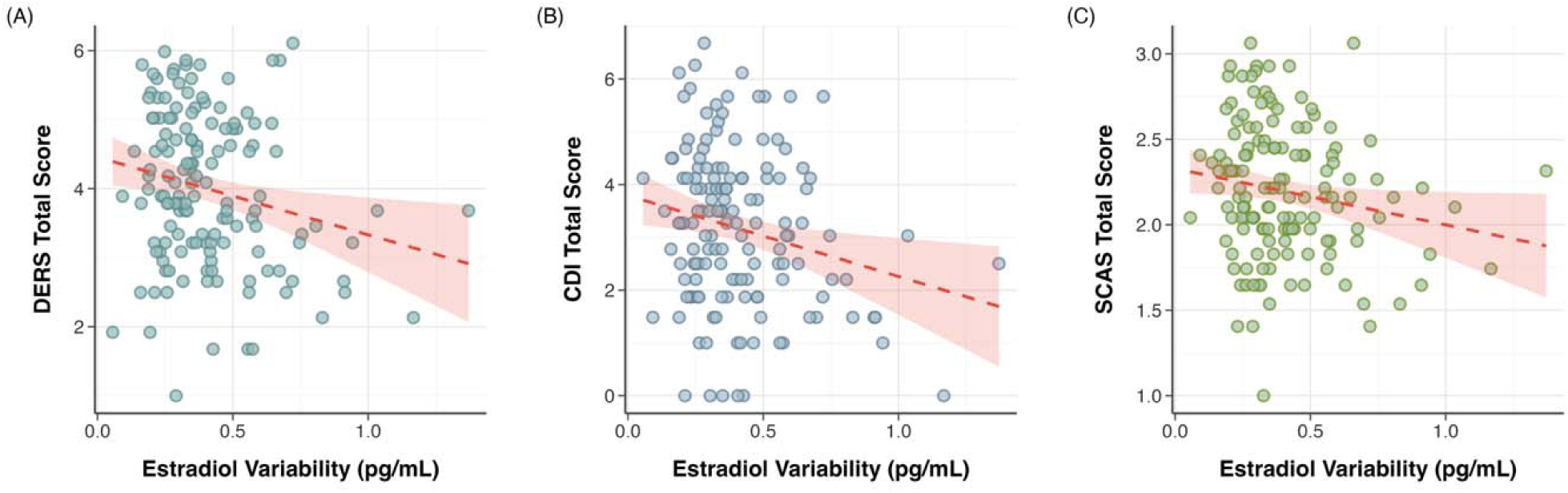
Negative association between E2 variability and A) DERS-SF score, B) CDI score, C) SCAS score. DERS-SF = Difficulties in Emotion Regulation Scale – Short Form; CDI = Children’s Depression Inventory; SCAS = Spence Children’s Anxiety Scale-Short. Dashed line represents the fitted linear regression with 95% CI (shaded area). Linear models were adjusted for menarche status, age at the time of MRI, adolescent’s race, primary caregiver’s education, and income-to-needs ratio

**Table 3:**
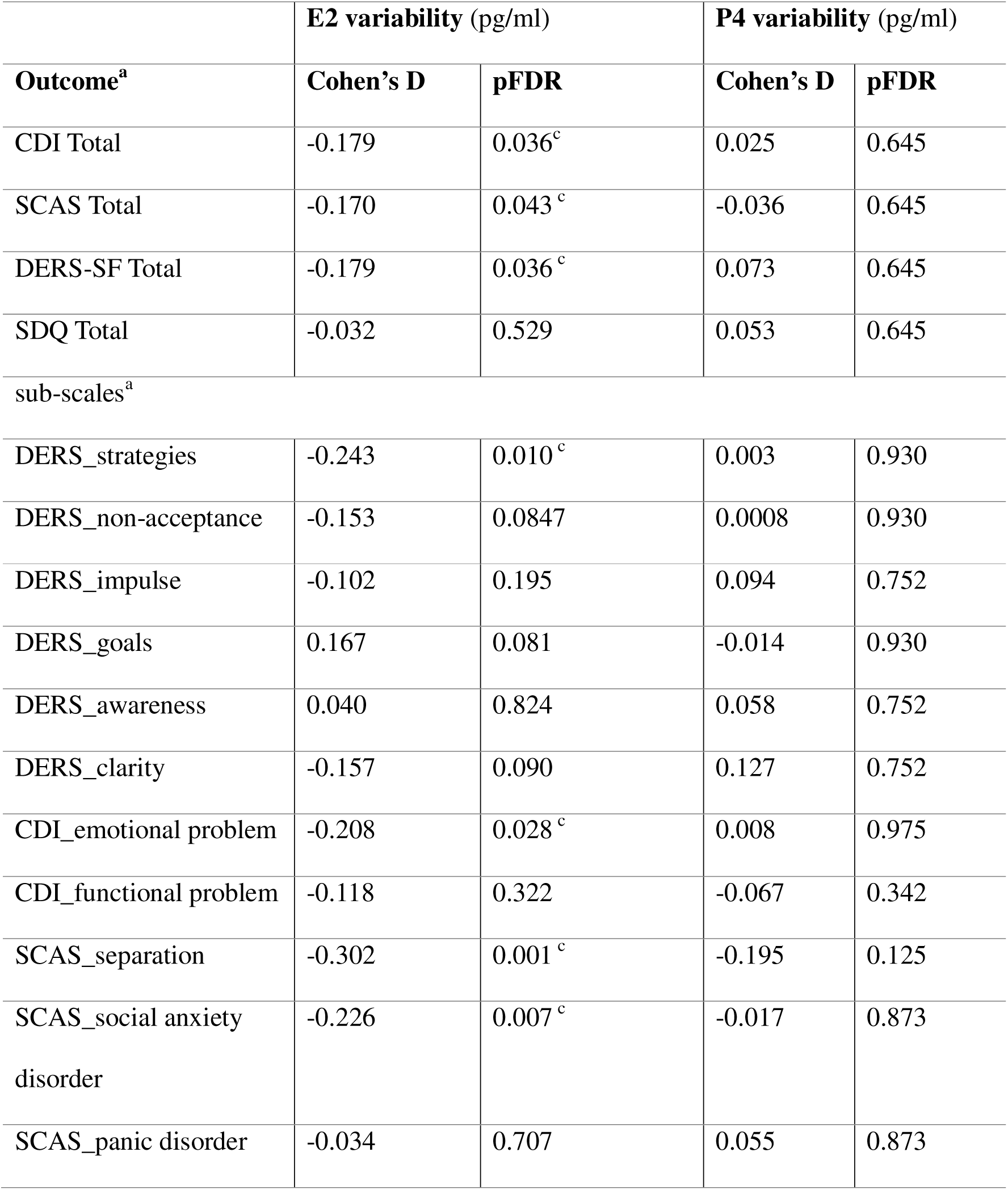
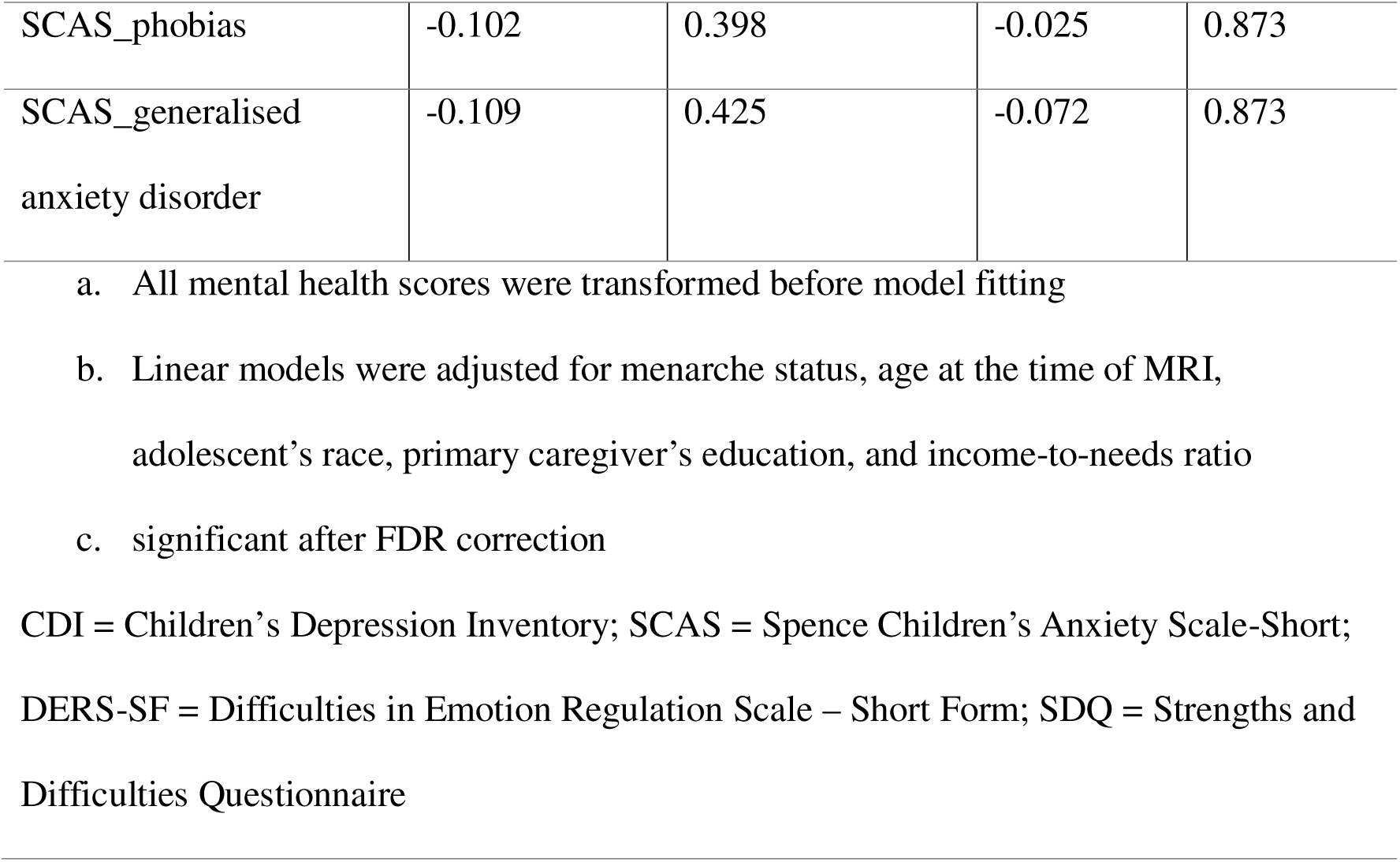
Linear regression output for models testing the association between hormone variability and mental illness symptoms/emotion regulation^b^.

### 3.3 Mediating role of Left Thalamus volume in the association between hormone variability and mental illness symptoms and emotion regulation

There were no significant indirect effects of P4 variability on mental health symptoms (CDI = −0.0005 [−0.0075 0.01], SCAS = −0.0001 [−0.0003 0.00], SDQ = −0.00074 [−0.00208 0.00]) or emotion dysregulation (DERS-SF = −0.0007 [−0.0022 0.00]) via left thalamus volume.

### 3.4 Exploratory analyses

#### i) Association between mean hormone levels and brain structure and mental illness symptoms/emotion regulation

Mean P4 levels were negatively associated with CDI total symptoms (Cohen’s d = −0.192, p = 0.022), but this effect did not survive multiple comparison correction (pFDR = 0.088). No other significant associations were found between mean E2 or P4 levels and brain structure or mental illness symptoms/emotion regulation. Refer to the SM Table 5 for model output.

#### ii) Menarche status or cycle irregularity as a moderator in the association between hormone variability and brain structure and mental illness symptoms/emotion regulation

Refer to SM for pre-versus post-menarche comparisons of hormone variability and correlation between Tanner stages and hormone variability There was a significant moderating effect of menarche status on the associations between E2 variability and right hippocampus volume, DERS total and CDI total scores. Simple slope analysis showed significant effects for the pre-menarche group only, with greater E2 variability associated with reduced right hippocampal volume, depressive symptoms and emotion dysregulation. Upon removal of detected influential outliers (as discussed in section 3.2 – I.A), the moderation effects became non-significant, but the effects in the pre-menarche group remained significant (Refer to SM for Figure 2 and model statistics). There were no significant moderating cycle irregularity effects on any of the mental illness symptoms, emotion dysregulation, or brain structure measures.

#### iii) Association between weekly hormone changes and weekly positive and negative affect

Linear mixed effect models showed a significant positive association between the absolute change in P4 levels from one week to the next and weekly negative affect scores in the corresponding week ( d = 0.115, p = 0.012). That is, greater week-to-week fluctuations in P4 levels were linked to increased negative affect. Additionally, there was a significant absolute change*direction effect in the association between weekly E2 levels and weekly positive affect ( d = 0.084, p = 0.048). A reduction in E2 levels from one week to the next was linked with a reduction in weekly positive affect (refer to SM).

## 4. Discussion

This study examined how hormone variability relates to brain structure and mental health problems in adolescent females. While we found that greater P4 variability was associated with lower left thalamus volume, we did not observe significant effects for other hypothesised regions. Additionally, we found that E2 variability was associated with lower depressive and anxiety symptoms, as well as emotion dysregulation. However, these associations were not mediated by brain structural changes as hypothesised.

Our finding that greater P4 variability was associated with smaller left thalamus volume may reflect a role of P4 in the maturation of the thalamus, a region with high expression of P4 receptors^43^. Prior longitudinal studies show that thalamic volume tends to decrease during early to mid-adolescence, to a greater extent in females compared to males^44^. Thus, our findings might suggest P4 variability as a biological mechanism partly responsible for thalamic development among females. Interestingly, structural changes in the thalamus have also been observed during other hormonal transitional stages, such as pregnancy and menopause^5^, as well as in menstrual cycle related clinical conditions such as PMDD, PMS and Primary Dysmenorrhea (PD), with studies reporting reduced thalamic volume associated with higher pain in women with these conditions^45^. Given the thalamus’s role in emotion regulation through its connections with the midbrain and cortex^46^, the current findings highlight it as a potential region through which P4 variability may influence emotional functioning during hormonal transition periods and disorders. However, we did not find links between P4 variability and mental health problems, directly or indirectly via thalamus volume. This null finding may indicate that P4-related changes in thalamus volume influence emotional (or other) functioning not measured in this study, or that effects on mental health may unfold over time.

Menarche or cycle regularity did not significantly moderate this effect, suggesting that the link between P4 variability and thalamic volume may be independent of menarche or the development of stable cyclic patterns. Further research is needed to clarify whether P4-related structural changes in the thalamus are specific to certain pubertal periods or persist into adulthood, which would help clarify whether these patterns reflect temporary developmental adaptations or stable individual differences in thalamic structure.

Contrary to our hypothesis, hormone variability was not associated with the structure of any other subcortical or cortical regions. Similarly prior study^25^ reported no vertex-wise associations, although they did find a negative relation between E2 variability and thickness in the right medial orbitofrontal cortex using an ROI approach. Notably, their study involved a smaller sample and a higher proportion of pre-menarche participants. Interestingly, in our exploratory subcortical ROI analyses, higher E2 variability was linked to smaller right hippocampus volume only among the pre-menarche group. Although speculative, this may reflect delayed brain maturation with higher E2 variability in this group, as these structures typically increase during adolescence^47^. However, future longitudinal studies are required to verify these patterns. Prior adolescent studies have linked average E2 or P4 levels to cortical density in prefrontal, parietal, and middle temporal cortices, and adult and animal studies have shown a reduced hippocampal volume during menstrual phases with average low E2 levels^23,24^. However, we found no associations for mean E2 or P4. This may partly reflect methodological differences, as our mean hormone estimates were based on four-five samples collected across a month, whereas prior studies used only two consecutive samples collected over two days. Additionally, previous work often included smaller female samples (<50 participants), did not account for menarchal status, and controlled for a specific cycle phase, all of which could contribute to inconsistent findings. Thus, future large-scale and replicable studies are needed to clarify hormone–brain associations.

In line with our hypothesised relationship between hormone variability and mental health problems, we found that increased E2 variability over a month was linked to reduced internalising symptoms and emotion regulation problems. To date, there are mixed findings regarding the direction of the relationship between E2 variability and mental health problems. For instance, a recent study has shown no association between E2 variability and depressive or anxiety symptoms in adolescent females, although their sample was relatively small and had limited variability in symptom levels^25^. In contrast, adult studies have shown that greater E2 variability during perimenopause was associated with heightened depressive symptoms^15,48^. Notably, our exploratory analysis indicated that the association between E2 variability and reduced mental health problems was stronger among pre-menarchal participants. This finding suggests that E2 variability during early adolescence may not have the same adverse implications observed during other hormonal transitions (e.g., postpartum, perimenopause). While speculative, greater E2 variability before menarche may reflect a normative feature of pubertal maturation or the gradual establishment of the biological mechanism, facilitating further emotional maturational processes. However, because symptom measures were collected concurrently with hormone sampling, and given the limited number of adolescent studies examining E2 variability, these interpretations should be considered preliminary and warrant cautious interpretation.

Interestingly, our within-person exploratory results indicated that a reduction in E2 levels from one week to the next was associated with less weekly positive affect, aligning with prior adolescent studies showing heightened mood sensitivity to E2 withdrawal and affective disturbances during other E2 withdrawal stages (i.e., peripartum symptoms)^4,49^. In addition, greater week-to-week change in P4 levels (regardless of the direction of change) was associated with greater weekly negative affect, suggesting that P4 instability may heighten sensitivity to negative mood. While the mechanisms by which weekly changes in E2 and P4 contribute to affect remain unclear, increased stress sensitivity during periods of hormonal fluctuation may be one potential pathway^50^. Moreover, heightened stress sensitivity has also been an important risk factor for the development of psychopathology during adolescence^51,52^. Thus, changes in weekly E2 and P4 levels that coincide with changes in affect may serve as early markers of emerging emotional patterns or vulnerabilities Notably, E2 changes showed a divergence in short-versus longer-term associations: while short-term reductions in E2 corresponded to lower positive mood, greater overall E2 variability across the month was associated with fewer mental health problems. This pattern suggests that transient hormonal declines may influence momentary mood states, whereas a more dynamic hormonal profile may reflect adaptive physiological responsivity. Further research is needed to clarify how greater hormonal variability, particularly when coupled with chronic or cumulative stress exposure, may contribute to the transition from normative affective sensitivity to later psychopathology.

## 5. Limitations and future directions

The current study is the first to examine how hormone variability relates to brain structure and mental health problems in adolescent females within a sample of this scale. However, several additional limitations should be considered when interpreting these findings.

Although the study captured weekly hormone data and self-reported menstrual cycle information, self-reports likely introduce error, particularly when adjusting for individual differences in menstrual cycle regularity among post-menarche girls. This limitation may have contributed to null associations when used as a moderator in exploratory analysis. Future research should aim to collect hormone samples at a higher frequency (e.g., every few days) and incorporate ovulation testing (e.g., using ovulation kits^53^) to more precisely map menstrual cycle phases and control for irregularity in cycle length.

Our study focused exclusively on E2 or P4. However, prior adolescent research has shown that other hormones, such as estrone (E1), demonstrate earlier increases during the pubertal transition compared with E2^4,54^. Future work should therefore examine a broader range of hormones to more fully characterise patterns of hormonal variability and their relevance for adolescent female development.

Further, Saliva was used as the collection medium because it is non-invasive and well-suited to frequent sampling in adolescents. Although plasma is the traditional gold standard for hormones like E2 and P4, saliva captures the free, biologically active fraction and may be more relevant for behaviour^55^. However, salivary assays are influenced by rapid fluctuations in hormone levels, necessitating frequent sampling for reliable estimates. While all biospecimens are affected by temporal hormone variation, saliva reflects these shifts more immediately, making sample timing and density particularly important for valid measurement^56^. Despite using a sensitive Salimetrics ELISA kit (lower limit: 0.998 pg/ml), four participants had values below the detection limit, indicating that approaches such as Liquid chromatography-tandem mass spectrometry (LC-MS/MS) and denser sampling schedules may be needed to improve accuracy for low-concentration hormone measurements^57^.

Finally, the current sample was recruited from Melbourne and comprised a community-based cohort, with a higher proportion of White/Caucasian participants (66.2%), and only a small number of individuals falling within the clinical range for mental illness symptoms. This limited diversity and restricted symptom range may have reduced the generalisability of our findings and contributed to the modest associations with mental illness symptoms (effect size ∼ 0.18). Replication in more diverse and clinically enriched samples will be important for determining the broader and clinical relevance of hormone–brain–behaviour associations.

## Conclusion

In conclusion, this study provides the first evidence of the links between E2 and P4 variability and brain structure and mental health problems in adolescent females. These findings lay an important foundation for future research examining how fluctuations in hormones contribute to brain development and risk for mental health difficulties across the pubertal transition. Moreover, the distinct associations observed between hormone variability, affect, and behaviour highlight the importance of investigating these dynamics in future developmental studies.

## CRediT authorship contribution statement

**MK:** Writing – original draft, Conceptualisation, Formal analysis, Visualisation. **YT:** Writing – review & editing, Conceptualisation, Supervision. **NV:** Conceptualisation, Supervision, Funding acquisition. **MH**: Writing – review & editing, Funding acquisition**. MC**: Writing – review & editing, Funding acquisition. **MS:** Writing – review & editing, Funding acquisition. **SW:** Conceptualisation, Methodology, Resources, Supervision, Project administration, Funding acquisition, Writing – review & editing.

