## Supplementary Material for "Oestradiol and Progesterone Variability in Adolescent Females: Associations with Brain Structure and Mental Health Problems"

### **2. METHODS:**

A total of 150 females participated in the study and provided weekly saliva samples over one month, while clinical and MRI assessments were conducted at a single point in time (refer to Figure 1 for the study design). Of these, only participants with a minimum of four saliva samples were included in the current analysis. Consequently, two participants with only three saliva samples were excluded. Furthermore, one participant with low-quality brain imaging data and two participants with exceptionally high E2 (> 15 pg/ml) and P4 (> 800 pg/ml) levels at one or more timepoints were also excluded. These hormone values were more than 20 and 9 standard deviations above the sample mean for E2 and P4, respectively, and exclusion of such values is consistent with prior research (Andersen et al., 2022). Thus, the final analytic sample comprised 145 participants. Of these, 27 provided four samples, 111 provided five, 6 provided six, and 1 provided seven samples. Notably, hormone variability did not differ by the number of samples. All participants were instructed to provide five weekly samples. However, the difference in saliva sample count primarily reflected missed weekly collections or rescheduling of the in-person visit.

#### 2.1 Magnetic resonance imaging (MRI)

**Data acquisition:**

- one participant with 0.8 mm and 192 slices, and ran for a smaller duration, ~5min. Other parameters; repetition time = 1900ms, echo time = 2.37ms, flip angle = 9
- six participants with 0.8mm, slices = 208, rt = 2100ms, echo time =2.51 ms, flip angle = 9

Reconstructed cortical surfaces were manually checked and rated out of 3 for quality purposes, such that participants with a rating of 1 (or low) were excluded from the analyses. Any issues with cortical segmentation (e.g., under/overestimation of white matter boundaries or dura mater included in pial surface) were manually corrected by adding control points or editing the voxels using FreeSurfer’s recon editing tools. The FreeSurfer recon pipeline was then re-run. In most cases, these corrections resolved the segmentation issues.

#### 2.2 Hormone (E2 and P4) variability measures and correlation:

**Other methods used to calculate hormone variability:**

1. Root Mean Square of Successive Differences
2. Mean Absolute Successive Difference (MASD)
3. Mean Squared Successive Difference (MSSD)

| **Table 1: Correlation between SD and other methods, MASD and MSSD** | | |
| --- | --- | --- |
|  | **E2 variability**  **r (p value)** | **P4 variability**  **r (p value)** |
| **MSSD** | 0.86 (<0.001) | 0.87 (<0.001) |
| **MASD** | 0.94 (<0.001) | 0.94 (<0.001 |

#### 2.3 Clinical and behavioural measures:

**SCAS-S:**

The Spence Children’s Anxiety Scale-Short (SCAS-S) is a 19-item self-report measure assessing the severity of anxiety symptoms in children and adolescents. It covers generalised anxiety disorder, separation anxiety disorder, social anxiety disorder, specific phobias and panic disorder. Each item scores from 0 (Never) to 3 (Always), with a maximum score of 57. The SCAS-S showed good internal consistency in our sample (Cronbach’s α = 0.86).

**CDI2:**

The Children’s Depression Inventory 2 (CDI-2) is a 28-item self-report measure assessing cognitive, affective, and behavioural symptoms of depression in children and adolescents. It includes two scales (Emotional Problems and Functional Problems) and four subscales: Negative Mood/Physical Symptoms, Negative Self-Esteem, Interpersonal Problems, and Ineffectiveness^1^. Items are scored from 0 to 2, with total scores ranging from 0 to 56; higher scores indicate more severe depressive symptoms. The CDI-2 demonstrated good internal consistency in this sample (Cronbach’s α = 0.87).

**SDQ:**

The parent-report Strengths and Difficulties Questionnaire (SDQ) was used to assess adolescents’ behavioural problems. This study focused on externalising symptoms, calculated as the sum of Conduct Problems and Hyperactivity/Inattention scores (5 items each, items are rated 0 (Not true), 1 (Somewhat true), or 2 (Certainly true). maximum score = 20). The SDQ externalising symptoms score showed acceptable internal consistency in this sample (Cronbach’s α = 0.72).

**DERS-SF:**

Difficulties in Emotion Regulation Scale – Short Form (DERS-SF) measures emotion regulation problems and is widely used in adults and adolescents. It has six sub-scales, each with 3 items: non-acceptance, difficulties engaging in goal-directed behaviour, impulse control difficulties, lack of emotional awareness, limited access to emotion regulation strategies, and lack of emotional clarity. Each item is scored from 1 to 5 (1 = Almost never to 5 = Almost always). Higher total scores indicate more difficulties in emotion regulation^2^. The DERS-SF has been validated in adolescents (ages 12-20) and has sound psychometric properties, including high correlation to the full-length version (subscales: .91 to .98). Internal consistency in the current sample was high (Cronbach’s α = 0.90).

2.4 PANAS:

The Positive and Negative Affect Schedule (PANAS) is a self-report measure of affect. It includes 20 items: 10 for positive affect (e.g., interested, excited, strong, enthusiastic, proud, alert, inspired, determined, attentive, active) and 10 for negative affect (e.g., stressed, upset, guilty, scared, hostile, irritable, ashamed, nervous, jittery, afraid). Items are rated from 1 (very slightly or not at all) to 5 (extremely), based on how participants felt over the past week. The PANAS has demonstrated good validity and internal consistency, with Cronbach’s α = 0.87 for positive affect and Cronbach’s α = 0.86 for negative affect in our sample.

#### 2.5 Exploratory Analyses:

i) To test whether any significant findings were specific to hormone variability, we also analysed the same linear regression models with mean E2 or mean P4 values as predictors instead of E2 or P4 variability. Mean E2 and mean P4 were calculated as the mean of all the available hormone measures within an individual.
ii) Additionally, we examined whether the effects of hormone variability on outcomes differed by menarcheal status and menstrual cycle regularity. Specifically, we conducted exploratory analyses testing whether menarche status (two groups: pre- and post-menarche) and cycle regularity (three groups: pre-menarche, regular, and irregular cycles) moderated the associations between hormone variability and brain structure, and mental illness symptoms. We also tested whether the indirect effect of hormone variability on mental health/emotion regulation via brain structure was moderated by menarche status or cycle irregularity.

***Classification method for regular and irregular menstrual cycles:***

Based on hormone levels and data from the past three cycles, we classified individuals into regular or irregular cycle categories. Specifically, those with an average cycle length over 45 days and maximum P4 levels below 50 pg/mL across all samples were grouped as irregular (or anovulatory) cycles. Individuals with maximum P4 levels of at least 100 pg/mL and maximum E2 levels of at least 1.5 pg/mL across all samples were classified as regular (or ovulatory) cycles. Participants (N = 9) whose P4 peaked between 50 and 100 pg/mL with E2 peak levels below 1.5 pg/mL were difficult to assign to either category and were therefore excluded. This left 139 participants for final analysis: 48 were premenarche, 28 postmenarche with regular cycles, and 63 postmenarche with irregular cycles.

iii) We additionally examined whether week-to-week changes in hormone levels were associated with weekly mood (positive and negative affect) changes. This would help us to understand whether hormone changes from one week to the next have some effects on concurrent measures of affect.

To investigate this, we calculated the difference in hormone levels from one week to the next (i.e., the value at each week minus the value from the previous week or (hormone levels)_n+1 week_ – (hormone levels)_n week_) within each individual. This allowed us to assess whether increases or decreases in hormone levels across a month were related to weekly changes in mood across a month. We ran two sets of linear mixed-effects models using the nlme package in R. In the first set, we tested whether absolute week-to-week changes in hormone (E2 or P4) levels were associated with weekly mood (positive or negative affect) outcomes. In the second set, we examined whether the direction of week-to-week change in hormone (increase vs. decrease) moderated this association. Each model included a nested random effect structure (~ 1 + Week | SubjectID) and the same covariates used in the main models (i.e., age, menarche status, child race, parent education, and income-to-needs ratio). To further probe significant interaction effects, we conducted simple slope analyses using the “sim_slopes” function in R.

### **3. RESULTS**

Group comparisons between pre- and post-menarche participants for hormone variability are in Table 1.

| Table 1: Group comparisons between pre- and post-menarche participants | | | |
| --- | --- | --- | --- |
|  | **Pre-menarche**  **Mean (sd)** | **Post-menarche**  **Mean (sd)** | **P value** |
| N | 46 | 99 |  |
| Age (years) | 12.34 (0.78) | 14.84 (1.52) | <0.001 |
| E2 variability (pg/ml) | 0.44 (0.27) | 0.39 (0.18) | 0.207 |
| P4 variability (pg/ml) | 48.33 (37.75) | 68.36 (43.11) | 0.008 |
| E2 mean (pg/ml) | 1.33 (0.67) | 1.66 (0.63) | 0.004 |
| P4 mean (pg/ml) | 113.82 (85.50) | 149.28 (98.13) | 0.037 |

| Table 2: Correlation between hormone variability and Tanner stages | | |
| --- | --- | --- |
|  | **E2 variability**  **r (p-value)** | **P4 variability**  **r (p-value)** |
| Tanner stage for breast development | 0.031 (0.718) | 0.186 (0.028) |
| Tanner stage for pubic hair development | 0.0171 (0.841) | 0.189 (0.025) |

| Table 3: Detailed breakdown of the specific reported races | |
| --- | --- |
| **Child_race** | **Count** |
| Asian | 22 |
| Asian,Other (please specify): | 1 |
| Asian,White/Caucasian | 7 |
| Indian | 6 |
| Indian,Other (please specify): | 1 |
| Indian,White/Caucasian | 2 |
| Other (please specify): | 6 |
| Pasifika/Maori,White/Caucasian | 2 |
| White/Caucasian | 96 |
| White/Caucasian,Other (please specify): | 2 |

| Table 4: Sub-scales description of mental health symptoms and emotion regulation | | |
| --- | --- | --- |
|  | **Min - Max** | **Mean (SD)** |
| SDQ_conduct | 0-5 | 0.81 (1.14) |
| SDQ_hyperactive | 0-8 | 1.99 (1.89) |
| DERS_Strate | 0-12 | 2.57 (2.45) |
| DERS_Non.ACC | 0-12 | 3.35 (2.97) |
| DERS_Impulse | 0-12 | 2.19 (2.68) |
| DERS_goals | 0-12 | 5.72 (3.20) |
| DERS_Awareness | 0-11 | 3.85 (2.50) |
| DERS_Clarity | 0-10 | 3.38 (2.68) |
| CDI_EmotionalProb | 0-17 | 4.55 (3.86) |
| CDI_FunctionalProb | 0-17 | 4.35 (3.22) |
| SCAS_SEP | 0-9 | 1.61 (1.75) |
| SCAS_SAD | 0-9 | 3.63 (2.12) |
| SCAS_PD | 0-9 | 2.02 (2.33) |
| SCAS_SP | 0-11 | 3.03 (2.15) |
| SCAS_GAD | 0-9 | 3.80 (2.25) |

#### 3.1 E2 variability and mental illness symptoms

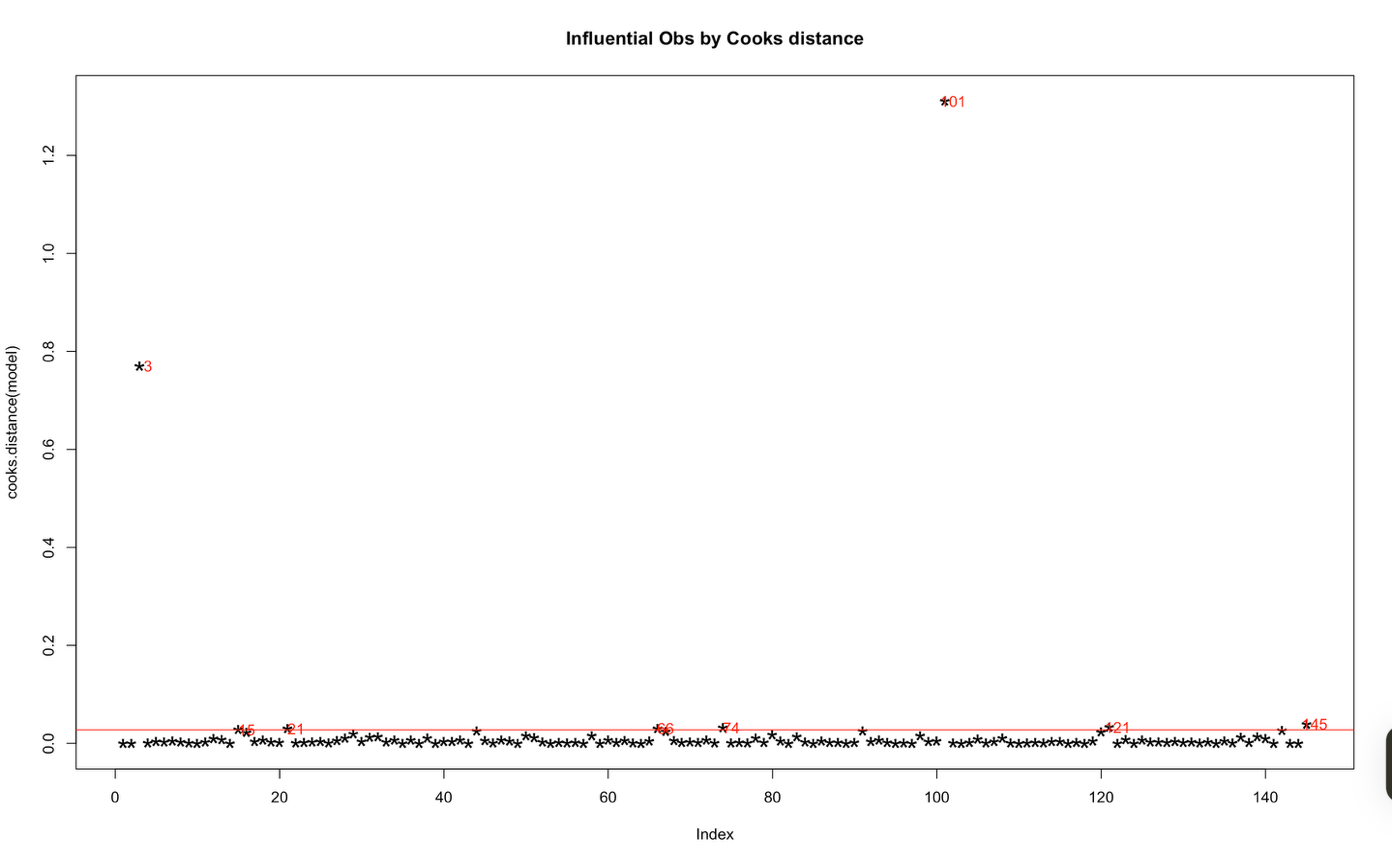

Figure 1: Influential outliers from the model between E2 variability and DERS total symptoms

| Table 4: Linear regression model between E2 variability and each of the measures with the full data^b^ | | |
| --- | --- | --- |
| **Behavioural measures^a^** | **Cohen’s D** | **pFDR** |
| CDI Total | -0.112 | 0.355 |
| SCAS Total | -0.123 | 0.355 |
| DERS Total | -0.038 | 0. .862 |
| SDQ Tot al | -0.002 | 0.976 |
| a. Transformed behavioural measures were used b. Linear models were adjusted for menarche status, age at the time of MRI, adolescent’s race, primary caregiver’s education, and income-to-needs ratio | | |

#### 3.2 Exploratory Analysis:

| Table 5: Linear regression model between mean hormone and the outcomes^b^ | | | | |
| --- | --- | --- | --- | --- |
|  | **E2 mean** | | **P4 mean** | |
| **Outcomes** | **Cohen’s D** | **pFDR** | **Cohen’s D** | **pFDR** |
| Left Thalamus | -0.197 | 0.266 | -0.138 | 0.884 |
| Right Thalamus | -0.057 | 0.699 | -0.062 | 0.884 |
| Left Caudate | 0.078 | 0.699 | -0.016 | 0.988 |
| Right Caudate | 0.045 | 0.699 | -0.067 | 0.884 |
| Left Putamen | -0.143 | 0.604 | -0.075 | 0.884 |
| Right Putamen | -0.075 | 0.699 | 0.007 | 0.997 |
| Left Pallidum | -0.044 | 0.699 | -0.050 | 0.884 |
| Right Pallidum | -0.033 | 0.745 | -0.047 | 0.884 |
| Left Hippocampus | -0.119 | 0.606 | -0.117 | 0.884 |
| Right Hippocampus | -0.063 | 0.699 | -0.051 | 0.884 |
| Left Amygdala | 0.006 | 0.933 | 0.028 | 0.935 |
| Right Amygdala | -0.051 | 0.699 | -0.0002 | 0.997 |
| Left Accumbens | -0.114 | 0.606 | -0.091 | 0.884 |
| Right Accumbens | -0.068 | 0.699 | -0.031 | 0.935 |
| **pFDR pFDR** | | | | |
| CDI Total | -0.003 | 0.965 | 0.009 | 0.904 |
| SCAS Total | -0.146 | 0.323 | -0.169 | 0.042 |
| DERS Total | -0.088 | 0.581 | -0.093 | 0.265 |
| SDQ Total | 0.0185 | 0.966 | 0.0432 | 0.603 |
| **pFDR pFDR** | | | | |
| CDI_EmotionalProb | -0.184 | 0.056 | -0.200 | 0.0346 |
| CDI_FunctionalProb | -0.078 | 0.348 | -0.111 | 0.182 |
| a. Transformed behavioural measures and sub-scales were used b. All Linear models were adjusted for menarche status, age at the time of MRI, adolescent’s race, primary caregiver’s education, and income-to-needs ratio;   For subcortical regions as the outcome, linear models were additionally adjusted for a dummy variable for different T1-w sequences, and ICV | | | | |

##### I. Menarche status as a moderator:

A. *E2 variability and the right Hippocampus volume moderated by menarche status*

SIMPLE SLOPES ANALYSIS:

Slope of sd_E2 when Menstruating_status = 0.00 (0):

Est. S.E. t val. p

--------- -------- -------- ------

-427.32	184.61	-2.31	0.02

Slope of sd_E2 when Menstruating_status = 1.00 (1):

Est. S.E. t val. p

-------- -------- -------- ------

219.67	184.72	1.19	0.24

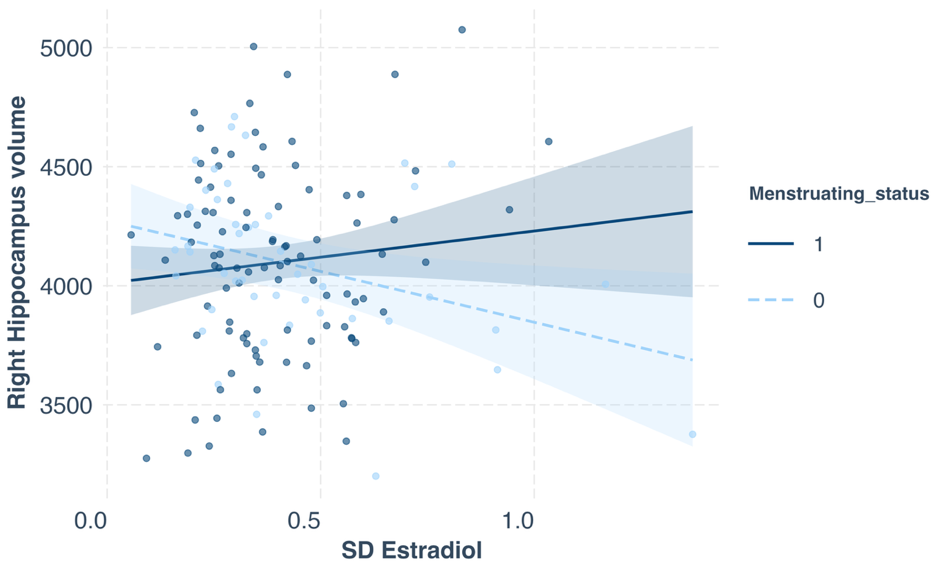

Figure 2: Represents the significant negative association between E2 variability and right hippocampal volume for the pre-menarche group (represented as “0” on the legend) only. Dashed line represents the fitted linear regression with 95% CI (shaded area). Linear models were adjusted for menarche status, age at the time of MRI, adolescent’s race, primary caregiver’s education, and income-to-needs ratio

B. *E2 variability and Mental illness measures moderated by menarche status – CDI and DERS – only for the measures that were significantly associated with E2 variability in the main analysis*

**CDI Total score ~ E2 variability**

SIMPLE SLOPES ANALYSIS

Slope of sd_E2 when Menstruating_status = 0.00 (0):

Est. S.E. t val. p

------- ------ -------- ------

-1.96	0.85	-2.31	0.02

Slope of sd_E2 when Menstruating_status = 1.00 (1):

Est. S.E. t val. p

------- ------ -------- ------

-0.64	0.88	-0.73	0.47

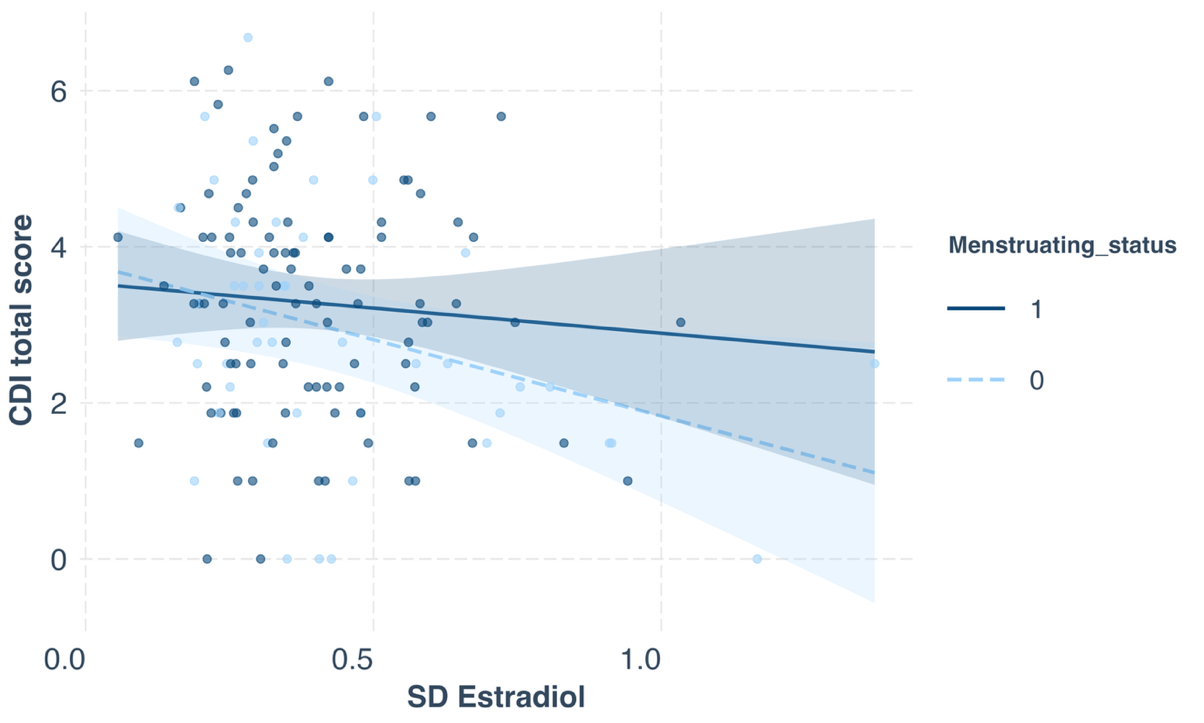

Figure 3: Represents the significant negative association between E2 variability and CDI total score for the pre-menarche group only (0). Dashed line represents the fitted linear regression with 95% CI (shaded area). Linear models were adjusted for menarche status, age at the time of MRI, adolescent’s race, primary caregiver’s education, and income-to-needs ratio

**DERS Total score ~ E2 variability**

SIMPLE SLOPES ANALYSIS

Slope of sd_E2 when Menstruating_status = 0.00 (0):

Est. S.E. t val. p

------- ------ -------- ------

-1.30	0.67	-1.94	0.05

Slope of sd_E2 when Menstruating_status = 1.00 (1):

Est. S.E. t val. p

------- ------ -------- ------

-0.76	0.69	-1.10	0.27

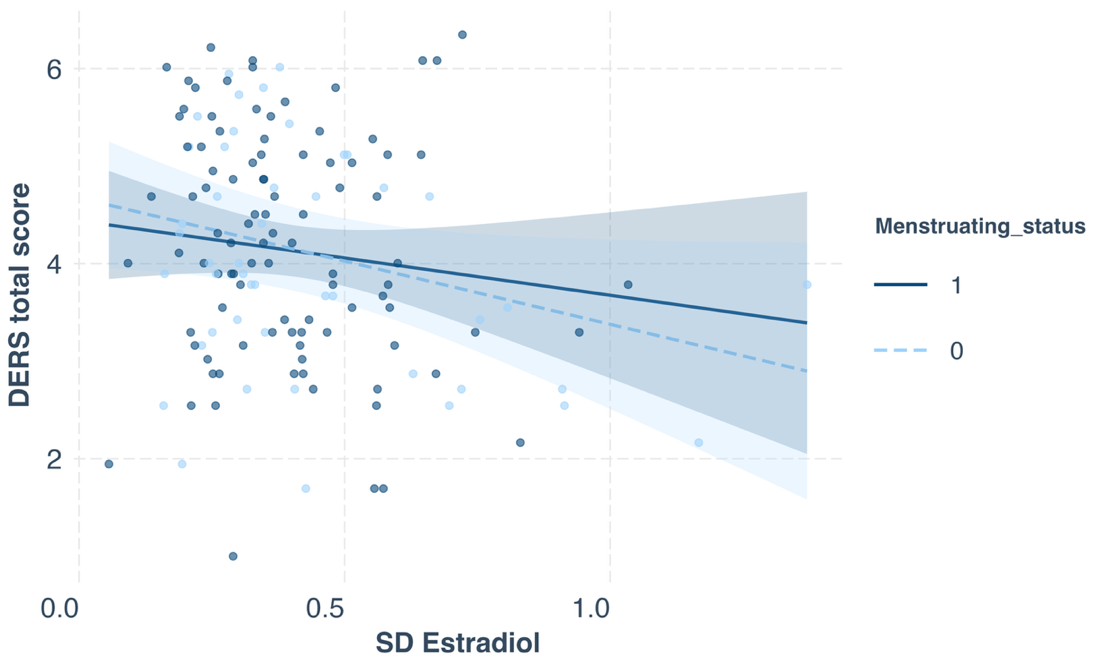

Figure 4: Represents the significant negative association between E2 variability and DERS total score for the pre-menarche group only (0). Dashed line represents the fitted linear regression with 95% CI (shaded area). Linear models were adjusted for menarche status, age at the time of MRI, adolescent’s race, primary caregiver’s education, and income-to-needs ratio

##### II. Association between the change in the hormones and mood affect:

**Weekly P4 change (absolute) and negative affect score**

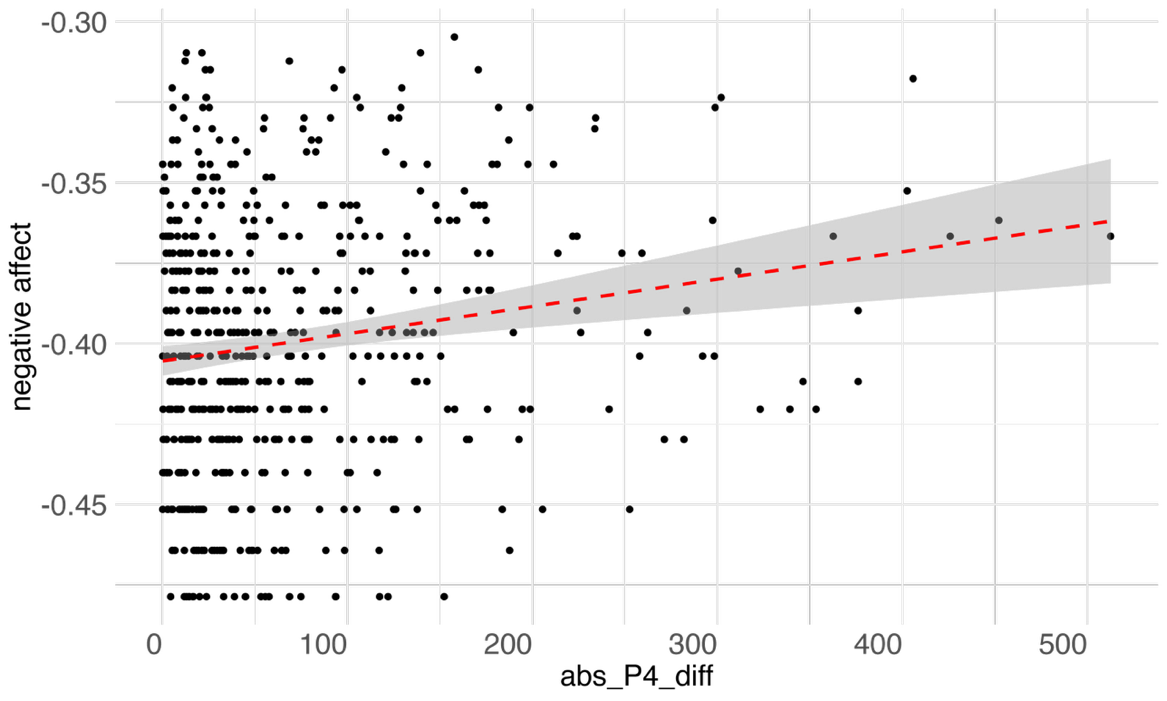

Figure 3: Represents the significant positive association between the absolute week-to-week change in P4 levels (pg/ml) and weekly negative affect score. Dashed line represents the fitted linear regression with 95% CI (shaded area). Linear models were adjusted for menarche status, age at the time of MRI, adolescent’s race, primary caregiver’s education, and income-to-needs ratio

**Weekly E2 change (with the direction of change) and positive affect**

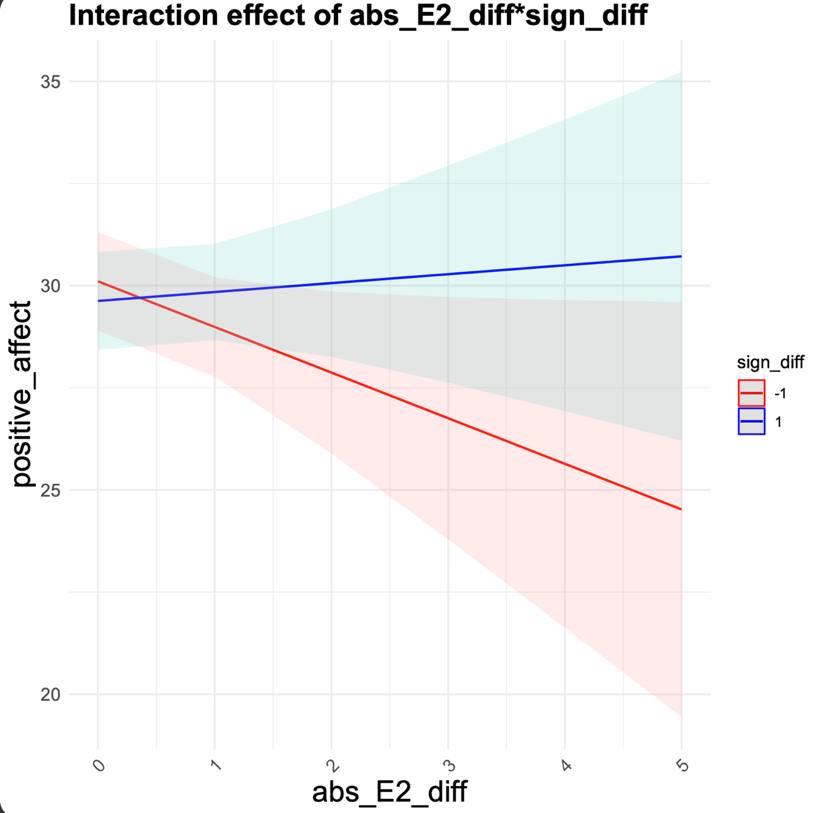

Figure 4: X-axis represents week-to-week difference in E2 levels (pg/ml). The red line represents the reduction in weekly E2 levels, whereas the blue line represents the increase in weekly E2 levels. The greater red slope shows that a reduction in weekly E2 levels significantly predicts a reduction in weekly positive affect score. Dashed line represents the fitted linear regression with 95% CI (shaded area). Linear models were adjusted for menarche status, age at the time of MRI, adolescent’s race, primary caregiver’s education, and income-to-needs ratio
